# A *TREM2*-derived circRNA is upregulated in the entorhinal cortex of Alzheimer’s disease patients

**DOI:** 10.1101/2021.06.23.449571

**Authors:** Amaya Urdánoz-Casado, Javier Sánchez-Ruiz de Gordoa, Maitane Robles, Miren Roldan, María Victoria Zelaya, Idoia Blanco-Luquin, Maite Mendioroz

**Author notes:** Corresponding author*: Maite Mendioroz, Mailing Address: NeuroEpigenetics Laboratory (Navarrabiomed-IdiSNA), c/Irunlarrea, 3 Pamplona-31008, Spain. Author e-mails:* Urdánoz-Casado A, Sánchez-Ruiz de Gordoa J, Robles M, Roldan M, Zelaya V, Blanco-Luquin I, Mendioroz M.

## Abstract

Circular RNAs (circRNAs) are a novel class of noncoding RNAs characterized by a covalent and stable closed loop structure. circRNAs are enriched in neural tissues, particularly at synapses, where they are involved in synaptic plasticity. Alzheimer’s disease (AD) is considered a synaptopathy since neurodegeneration causes loss or dysfunction of synapses. Microglia participate in synaptic pruning and also play a crucial role in developing AD. For instance, genetic variants in *TREM2*, a microglia-related gene, are risk factors for AD. Alterations in circRNAs expression have been described in different neurological diseases, including AD. However, no *TREM2*-derived circRNAs have been described so far. *TREM2* has 3 linear RNA variants due to alternative splicing. We hypothesized that alternative splicing of exon 4 might be favoring circRNAs originating from *TREM2* (circTREM2s), which in turn might be involved in AD pathogenesis. First, divergent primers (overlapping exons 3-4 and 4-5) were designed to amplify circRNAs by RT-PCR, which were confirmed by Sanger sequencing. Three candidate *TREM2*-derived circRNAs were identified on control human entorhinal samples. Then, additional primer sets were used to confirm back-splicing junctions. One of the circRNAs, circTREM2_1, was consistently amplified with all primer sets. In addition, circTREM2_1 was also present in AD entorhinal cortex samples and in HMC3 cells. We observed that circTREM2_1 is up-regulated in AD entorhinal cortex samples compared to controls, particularly at early stages of the disease, when we performed RT-qPCR. In conclusion, we have identified a novel circRNA derived from the *TREM2* gene that could play a role in AD pathogenesis.

## Introduction

Circular RNAs (circRNAs) are a novel class of non-coding RNA characterized by the covalent junction between the 3’ end and 5’end, generated by back-splicing process. From the same pre-mRNA precursor, messenger RNAs (mRNA) and circRNAs are formed by canonical splicing and alternative splicing (1–4). In the circRNAs biogenesis not only the spliceosome machinery and the RNA polymerase II are implicated but also other elements, like *Alu* elements, ADAR (adenosine deaminase that acts on RNA) enzyme or RBP (RNA binding proteins) which participate in the process (1, 5–10). This covalent union gives circRNAs a more stable structure than linear RNAs, since they are resistant to the action of RNases (1), and this characteristic, among others, makes circRNAs interesting molecules to investigate in human diseases.

Moreover, circRNAs are highly evolutionary conserved, and their expression has been observed across a wide range of body tissues (1, 4, 10, 11). However, it is in the brain where they are most expressed. circRNAs are enriched in neuronal tissues, in fact it has been reported that 20% of brain genes generate circRNAs (12, 13). Interestingly, neuronal specialization is related to a high level of alternative splicing (14) and this process could also explain the huge amount of brain-specific circRNAs (10).

Nevertheless, circRNAs expression levels vary from one brain structure to another, being synapsis the place where their expression is highest (10, 12, 13). Their expression is also modified along neuronal development, changing space-timely and increasing its synthesis (10). All these data suggest that RNA circularization is crucial for neuronal function. Alterations have been described in circRNAs expression levels in different neurological diseases, like multiple sclerosis, Parkinson’s disease, amyotrophic lateral sclerosis, and Alzheimer’s disease (AD) among others (15–18).

AD is a neurodegenerative disease that is the leading cause of age-related dementia (19). In terms of anatomopathological changes, it is characterized by extracellular accumulation of amyloid plaques and intraneuronal tau neurofibrillary tangles along with loss of synapses between neurons (20–22). At present, AD is considered a complex disease, probably caused by the interaction between 3 different factors: genetic, environmental and epigenetic (20–22). To date, the most widely studied epigenetic mechanisms are DNA methylation and posttranscriptional histone modifications (23, 24). However, it has been described that non-coding RNAs, such as circRNAs, also play an important role in epigenetic regulation (25, 26). The best known circRNA implicated in AD is CDR1as (ciRS-7), generated from Cerebellar Degeneration Related Protein 1 (*CDR1*) gene, which has more than 70 union sites for microRNA-7 (miR-7) (4, 27, 28). In AD, differences of CDR1as expression levels have been observed in the hippocampus (29, 30), one of the most vulnerable brain regions to tau deposition. In addition, *APP* and *MAPT* are genes encoding the main proteins involved in AD, i.e. amyloid precursor protein and tau protein, which circRNAs are also transcribed from (31, 32). Most interestingly, Aβ peptides may be synthesized from an APP-derived circRNA, known as circAβ-a (32, 33). Recent studies in AD-affected brain regions have also found differential expression in a number of circRNAs like circHOMER1 and circCORO1C (18, 34, 35).

On the other hand, microglia, the innate immune cells of the brain, are also implicated in the AD pathogenesis (36, 37). Over the past few years, the role of microglia in AD pathogenesis has radically changed, gaining a more relevant position in neurodegenerative diseases. In this scenario, microglia-related genes are receiving special attention, as is the case with *TREM2* gene.

*Trem2* (Triggering receptor expressed on myeloid cells 2) is a cell surface protein almost exclusive of microglia cells and tightly associated with AD. Some single nucleotide polymorphisms (SNPs) in this gene have been described as risk factors of AD, like p.R47H (rs75932628) or p.R62H (rs143332484) (38, 39). Moreover, in cerebrospinal fluid (CSF) of AD patients the soluble form of *Trem2* (sTREM2) is increased (39). sTREM2 is thought to be originated by action of proteases on the extracellular portion of *Trem2* (C-terminal fragment)(40). Thus far, 3 distinct *TREM2* transcripts have been described expressed in the brain, i.e. ENST00000373113, ENST00000373122 and ENST00000338469 (38, 41). The first transcript (ENST00000373113) consists of 5 exons representing the canonical and longest *TREM2* isoform, the second transcript (ENST00000373122) is shorter than the canonical one due to the lack of exon 5 and part of exon 4, meanwhile the third transcript (ENST00000338469) is the result of the alternative splicing of exon 4, which encodes the transmembrane domain; therefore, sTREM2 may also be translated from the latter transcript (Supplemental Figure 1). Both first and third isoforms have been reported to be elevated in AD brain tissues (38, 41). Interestingly, bioinformatics by using the Genome Browser *SIB Alt-Splicing Track Settings (42)* also predicts that alternative splicing may involve exon 3, 4 and 5. Indeed, Yanaizu et al. described that exon 3, under normal conditions, undergoes alternative splicing in order to regulate *TREM2* expression (43). Also, it has been noted an increased expression of exon 3 and 4 depending on the degree of AD-related neurofibrillary pathology (44). It is also interesting to note that epigenetic mechanisms involving *TREM2* are also modulated in AD, since *TREM2* DNA methylation and hydroxymethylation levels are known to be altered in the hippocampus (45)and other brain regions affected by AD (46–48).

Despite the increasingly relevant involvement of circRNAs in the brain and the key role of *TREM2* in neurodegenerative processes, no circRNAs originating from *TREM2* have been described in AD so far. After examining the genetic structure of *TREM2*, we hypothesize that alternative splicing of exon 4 could be favoring *TREM2*-originated circRNAs (circTREM2s), which in turn may be involved in the pathogenesis of the AD. We decided to study exon 4 as this exon encodes the transmembrane domain and is alternatively spliced to form the sTREM2 isoform. Thus, we used a Human embryonic microglia cell line (HMC3 cells) and human entorhinal cortex, a brain region most vulnerable to AD pathology, from AD patients and controls to identify *TREM2* originated circRNAs.

## Material and Methods

### Human Entorhinal Cortex Samples

Brain entorhinal cortex samples from 27 AD patients and 16 controls were provided by Navarrabiomed Brain Bank. After death, half brain specimens from donors were cryopreserved at − 80 °C. Neuropathological examination was completed following the usual recommendations (49) and according to the updated National Institute on Aging-Alzheimer’s Association guidelines (50).

Assessment of Aβ deposition was carried out by immunohistochemical staining of paraffin-embedded sections (3–5 μm-thick) with a mouse monoclonal (S6 F/3D) anti-Aβ antibody (Leica Biosystems Newcastle Ltd, Newcastle upon Tyne, United Kingdom). Evaluation of neurofibrillary pathology was performed with a mouse monoclonal antibody anti-human PHF-TAU, clone AT-8 (Tau AT8) (Innogenetics, Gent, Belgium), which identifies hyperphosphorylated tau (p-tau) (51). The reaction product was visualized using an automated slide immunostainer (Leica Bond Max) with Bond Polymer Refine Detection (Leica Biosystems, Newcastle Ltd). Other protein deposits, such as synuclein deposits, were ruled out by a monoclonal antibody against α-synuclein (NCL-L-ASYN; Leica Biosystems, Wetzlar, Germany). The staging of AD was performed by using the ABC score according to the updated National Institute on Aging-Alzheimer’s Association guidelines (50). ABC score combines histopathologic assessments of Aβ deposits determined by the method of Thal (A) (50), staging of neurofibrillary tangles by Braak and Braak classification (B) (51), and scoring of neuritic plaques by the method of CERAD (Consortium to Establish A Registry for Alzheimer’s Disease) (C) (52) to characterize AD neuropathological changes. Thus, the ABC score shows three levels of AD neuropathological severity: low, intermediate, and high. A summary of the characteristics of subjects considered in this study is shown in Supplemental Table 1.

### Cell Culture

Human embryonic microglia clone 3 (HMC3, ATCC^®^ CRL3304™) cells were cultured in minimum essential medium (MEM) Eagle-EBSS with NEAA without L-Glutamine (Lonza, Cologne, Germany) supplemented with 10% fetal bovine serum (Sigma-Aldrich, Saint Louis, MO, USA), 1% penicillin-streptomycin (Gibco™ Thermo Fisher Scientific, Waltham, MA, USA,) and 1% L-glutamine 200mM (Gibco™, Thermo Fisher Scientific, Waltham, MA, USA) at 37°C and in an atmosphere of 5% CO2.

### RNA isolation and Reverse transcription – polymerase chain reaction (RT-PCR)

Total RNA including small RNA species was isolated from cells and entorhinal cortex samples with miRNAeasy mini Kit (QIAGEN, Redwood City, CA, USA) following the manufacturer’s instructions. Concentration and purity of RNA were both evaluated with NanoDrop spectrophotometer. Complementary DNA (cDNA) was reverse transcribed from 500 ng total RNA with SuperScript^®^ III First-Strand Synthesis Reverse Transcriptase (Invitrogen, Carlsbad, CA, USA) after priming with random primers. RT-PCR was performed by using GoTaq^®^ DNA polymerase (Promega, 2800 Woods Hollow Road·Madison, USA) in an Applied Biosystems™ Veriti™ Thermal Cycler, 96-Well (Applied Biosystems, Foster City, CA, USA). PCR conditions were as follows: denaturation at 95°C for 20 s, extension at 72°C for 30 s and annealing temperatures 60°C for 40s and cycles used were 40. Primer3 software was used for divergent primers design (Supplemental Table 2).

### Candidate band selection and Sanger sequencing

Candidate bands were selected after 1.8% agarose gel electrophoresis of RT-PCR products. Bands purification were made with Wizard^®^ SV Gel and PCR Clean-Up System (Promega, 2800 Woods Hollow Road·Madison, USA). Next, Sanger sequencing was performed and UCSC (University of California Santa Cruz) Genome Browser software was used for the sequence alignment (42).

### Real time quantitative PCR (RT-qPCR) assay

Total RNA were isolated from entorhinal cortex with RNAeasy Lipid Tissue mini Kit (QIAGEN, Redwood City, CA, USA) following the manufacturer’s instructions. Genomic DNA was removed with recombinant DNase (TURBO DNA-free™ Kit, Ambion). Concentration and purity of RNA were both evaluated with NanoDrop spectrophotometer. cDNA was reverse transcribed from 500 ng total RNA with SuperScript^®^ III First-Strand Synthesis Reverse Transcriptase (Invitrogen, Carlsbad, CA, USA) after priming with random primers. RT-qPCR reactions were performed in triplicate with TaqMan Master Mix: Premix Ex Taq (TaKaRa, Otsu, Japan) in a QuantStudio 12K Flex Real-Time PCR System (Applied Biosystems, Foster City, CA, USA). Sequences of primer pair and probe were designed using Real Time PCR tool (IDT, Coralville, IA, USA) (Supplemental Table 2). Relative expression level of mRNA in a particular sample was calculated as previously described (53) and *ACTB* gene was used as the reference gene to normalize expression values.

### Quantitative assessment of Aβ deposits in brain tissues

In order to quantitatively assess the Aβ burden for further statistical analysis, we applied a method to quantify protein deposits. This method generates a numeric measurement that represents the extent of Aβ deposition. Sections of the entorhinal cortex were examined after performing immunostaining with anti Aβ antibody as described above in Human Entorhinal Samples. Three pictures were obtained for each immunostained section by using an Olympus BX51 microscope at ×10 magnification power. Focal deposit of Aβ, as described by Braak & Braak (neuritic, immature, and compact plaque) (51), was manually determined and was further edited and analysed with the ImageJ software. Then, Aβ plaque count, referred to as amyloid plaque score (APS) and total area of Aβ deposition was automatically measured by ImageJ and averaged for each section.

### Statistical Analysis

Statistical analysis was performed with SPSS 25.0 (IBM, Inc., USA). Before performing differential analysis, we checked whether continuous variables follow a normal distribution, as per one-sample Kolgomorov-Smirnov test and the normal quantil-quantil (QQ) plots. The values of expression levels were transformed by using log10(X) to meet the conditions of normality, then T-student test was used to analyze differences in the expression levels of circTREM2_1. Significance level was set at p-value < 0.05. One-way analysis of variance (ANOVA) and *posthoc* Bonferroni test was used to analyze differences in the expression levels of circTREM2_1 among ABC stage groups. Pearson test was used to assess correlation between continuous variables. GraphPad Prism version 6.00 for Windows (GraphPad Software, La Jolla, CA, USA) was used to draw graphs.

## Results

### circTREM2 identification in entorhinal human brain

So far, no circTREM2 has been described. *TREM2* has 3 isoforms due to alternative splicing (38, 41) and alternative splicing of exon 3 and exon 4 could have some implication in AD pathology (43, 44). So, we hypothesize that exon 4, which is skipped to form sTREM2, may be involved in the circRNA biogenesis from *TREM2*.

First, we designed divergent primers circTREM2_3-4 and circTREM2_4-5 (Figure 1A, Supplemental Table 2), in order to amplify circular, but not linear, RNA template that includes exon 4. Divergent primers are head-to-head oriented primers that target the exon or exons of interest. Next, we performed RT-PCR in RNA isolated from human entorhinal cortex without neuropathological changes of any neurodegenerative process, known as control samples. circTREM2_3-4 primers succeeded to amplify a range of different molecular weight PCR products; in contrast, circTREM2_4-5 primers did not. After electrophoresis, we selected a number of PCR products from agarose gels for Sanger sequencing analysis (Figure 1B, Supplemental Figure 2 and Supplemental Table 2). Following this approach, we identified 3 different sequences that included candidate junctions between, on the one hand, exons 2 and 5 (circTREM2_1, circTREM2_2) and, on the other hand, intron 4 and exon 2 (circTREM2_3) (Figure 1A, B, C, D, Supplemental Figure 2 and Supplemental Table 2) in the human entorhinal cortex. As expected, the order of exons found in the sequences (Figure 1 C, D) did not correspond to the RNA linear form. Therefore, these sequences are assumed to correspond to circRNAs and include circRNAs back-splicing junctions.

**Figure 1:**
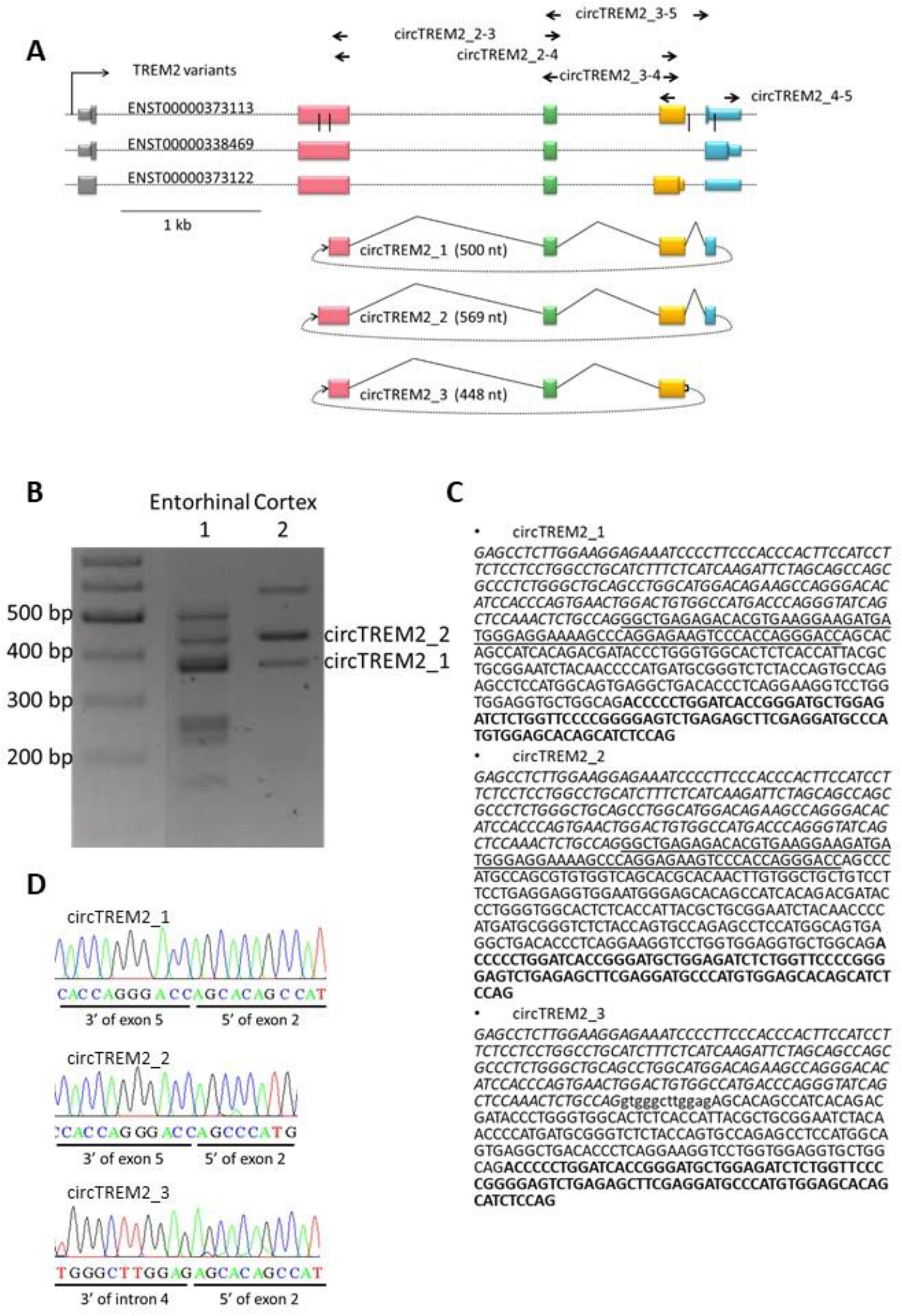
*TREM2* and circTREM2 representation. A) Gene map of linear and circular *TREM2* RNA. Boxes represent exons, black vertical lines are back-splicing junction sites. Dotted lines indicated the back splicing of exons. Arrows represent set of primers used. B) Representative figure of RT-PCR products of circTREM2_3-4 primer set in agarose gel. C) Sequences of circTREM2s, circRNA part underlined correspond to exon 5, regular letters to exon 2, italic letter to exon 4, letter in bold to exon 3, and lower case to intron. D) Electropherograms show the sequence of the junction of each circTREM2, confirming the fusion between 3’ of exon 5 and 5’ of exon 2 in the case of circTREM2_1 and circTREM2_2, and the fusion between 3’ of intron 4 and 5’ of exon 2.

To make sure these sequences are part of circRNAs, we designed additional divergent primers but, in this case, overlapping other *TREM2* exons, namely circTREM2_2-4, circTREM2_2-3, circTREM2_3-5 (Supplemental Table 2). After RT-PCR and Sanger sequencing, we observed that one of the three sequences described above, circTREM2_1, was amplified by using all 3 different primer sets (circTREM2_2-4, circTREM2_2-3 circTREM2_3-5). circTREM2_1 seems to correspond to a circRNA formed by exons 2, 3, 4 and 5, whose circRNA junction is located between part of exon 2 and exon 5 (Figure 1 A, C and D). Regarding the other two sequences (circTREM2_2 and circTREM2_3) (Figure 1 A, C and D), they probably may be described as circRNAs, but were not able to be detected with all primer sets.

To be noted, circTREM2_2 includes a SNP describe as a risk factor for AD (rs143332484) and the 3’end of circTREM2_1 and circTREM2_3, when aligned to the human genome, is only 55 bp distant from rs143332484 and 100 bp away from rs75932628 (Supplemental Figure 3), both genetic variants associated with AD.

Once we identified circRNAs derived from the *TREM2* gene in controls, we decided to perform the same approach in the entorhinal cortex of AD samples. In this case, we obtained similar results, circTREM2_1 was the circRNA that we detected with the highest consistency.

### circTREM2 in human microglia cells

In brain, *TREM2* is a gene that is almost exclusively expressed in microglia. Microglia involvement in AD is being extensively investigated, although its role in the disease is still unclear, both protective and deleterious roles have been attributed to it (54). Thus, we inquired whether any of the circTREM2 described above were expressed in microglia.

To address this question, we performed RT-PCR on RNA isolated from HMC3 cells, a human embryonic microglia cell line. We were able to confirm that both circTREM2_1 and circTREM2_3 were expressed in human microglia, although circTREM2_3, as in the case of human entorhinal cortex, was not detected with all primer sets.

### circTREM2 differential expression in AD entorhinal cortex

In order to measure the expression levels of circTREM2 and to know whether a differential expression was observed between AD patients and controls, we carried out a RT-qPCR assay. All samples passed the RNA quality threshold, so 27 AD samples and 16 controls from entorhinal cortex were studied. For this experiment, we selected circTREM2_1, since it was the one most consistently detected in our samples. To ensure that only circTREM2_1 was amplifying in the RT-qPCR assay, we designed a specific TaqMan-probe overlapping the back-splicing junction (Supplemental Table 2). Thus, we observed that circTREM2_1 was upregulated in AD samples (Fold-change = 2.53, p-value <0.05) compared to controls (Figure 2 A). When we examined the expression levels of circTREM2_1 considering the ABC score, we found that expression in the low ABC score cases was significantly increased compared to controls (p-value<0.05) (Figure 2 B).

**Figure 2:**
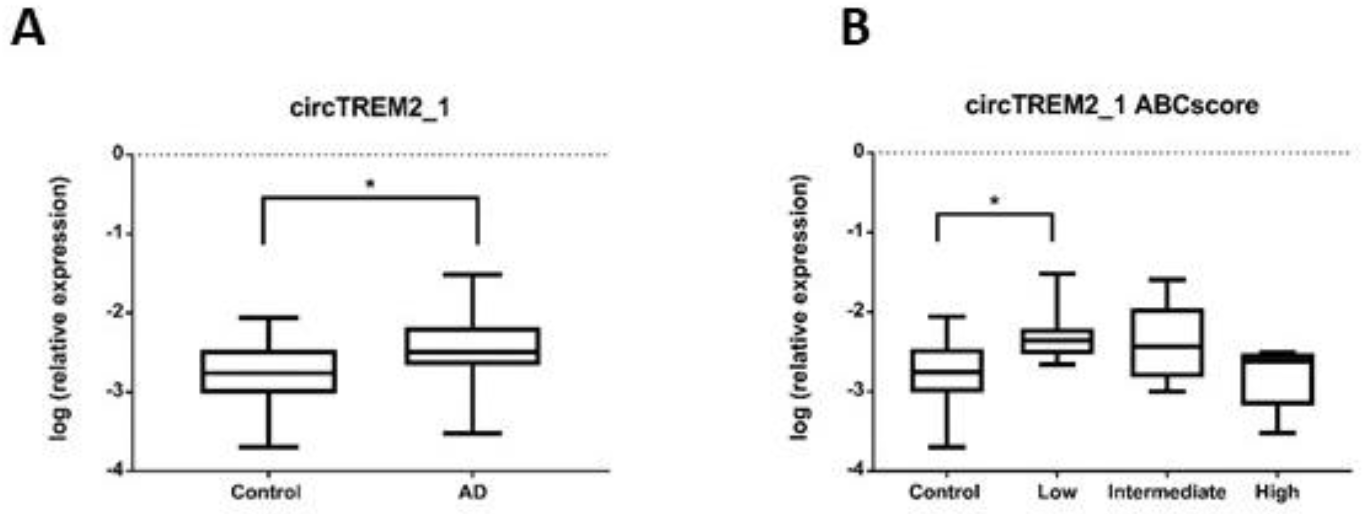
circTREM2_1 expression levels relative to ACTB housekeeping gene expression. A) The graph shows a significant increase in circTREM2_1 levels in AD entorhinal cortex samples compared to Control entorhinal cortex samples. B) circTREM2_1 expression decreased across AD stages, as shown when circTREM2_1 expression levels are sorted by ABC score. Vertical lines represent the standard error of the mean.*p-value<0.05.

### circTREM2 and Aβ deposits

TREM2 has been described as having an affinity for Aβ peptide, and this affinity is implicated in the activation of the signaling cascade for the clearance of Aβ deposits in AD (55–57). Therefore, we wondered if circTREM2_1 expression showed any correlation with Aβ burden. We observed a moderate negative correlation between global average area of Aβ deposits in entorhinal cortex and circTREM2_1 expression level among the AD cases (r=−0.434, p-value=0.05).

### Prediction of miRNAs binding sites within circTREM2s

In order to assess the functional role of circTREM2, we used bioinformatics tools to predict miRNAs binding sites within the circRNAs sequences. The miRDB software (58, 59) was employed to find associated miRNAs. A total of 16 miRNAs was predicted to join with circTREM2_1, circTREM2_2 or circTREM2_3 (Table 1). Some of these miRNAs have been described to be involved in human pathological processes. For instance, miR-765 is implicated in osteogenesis (60) and has been described as a tumor suppressor (61). miR-939 seems to inhibit apoptosis and regulate angiogenesis through the nitric oxide signaling pathway mainly in the context of coronary disease (62). However, no circTREM2-related miRNAs have been associated with neurodegeneration or brain diseases so far.

**Table 1:** miRNAs predicted with miRBD sofware to bind circTREM2s.

| miRNAs predicted to join | circTREM2_1 | circTREM2_2 | circTREM2_3 |
| --- | --- | --- | --- |
| hsa-miR-765 | Yes | Yes | No |
| hsa-miR-11181-3p | Yes | Yes | No |
| hsa-miR-6890-3p | Yes | Yes | Yes |
| hsa-miR-6766-5p | Yes | Yes | No |
| hsa-miR-6756-5p | Yes | Yes | No |
| hsa-miR-653-3p | Yes | Yes | Yes |
| hsa-miR-6770-5p | Yes | Yes | No |
| hsa-miR-6131 | Yes | No | No |
| hsa-miR-6762-3p | Yes | Yes | No |
| hsa-miR-4783-3p | Yes | Yes | Yes |
| hsa-miR-2392 | Yes | Yes | No |
| hsa-miR-4483 | Yes | Yes | No |
| hsa-miR-6745 | Yes | Yes | No |
| hsa-miR-6749-3p | No | No | Yes |
| hsa-miR-939-3p | No | No | Yes |
| hsa-miR-3657 | No | No | Yes |

## Discussion

In this study, a *TREM2*-derived circRNA transcript (circTREM2_1) has been consistently identified in the human brain for the first time. We also show evidence of other circTREM2s in the human brain. In addition, we observed that circTREM2_1 is expressed in a cell model of human microglia. Interestingly, an increase in expression levels of circTREM2_1 was found in the entorhinal cortex, a brain region most vulnerable to AD pathology, between AD patients and controls. This increase was shown mainly at early stages of AD and negatively correlated with Aβ deposits in the entorhinal cortex.

*TREM2* gene encodes a transmembrane glycoprotein receptor crucial in regulating microglia function. Microglia are the innate immune cells in the brain and usually perform relevant surveillance tasks. In microglia, *TREM2* promotes phagocytosis of apoptotic cells and debris, misfolded proteins and also decreases cytokines production, which is therefore of great relevance to maintain brain homeostasis (63–65). Though, this homeostasis is altered in brain diseases, such as neurodegenerative conditions. Some microglial cells are then transformed into a particular state, known as disease-associated microglia (DAM)(66). DAM was first described as a subset of brain immune cells characterized by a specific molecular signature identified by single-cell transcriptomics performed on brain immune cell samples of neurodegenerative diseases (66). It has also been reported that DAM may early sense neuronal damage and play a protective role in neurodegeneration (67). DAM is mainly detected in brain regions affected by AD and colocalize with Aβ plaques (67). Interestingly, microglia seem to use a molecular mechanism, which involve the Trem2 signalling pathway, to sense and response to brain damage (67). *TREM2* gene is crucial to activating DAM in a neurodegenerative environment (66, 68) since *TREM2* senses a wide range of damage-signals and is able to maintain the DAM phenotype in response to these cues (67).

In this scenario, regulation of *TREM2* transcripts expression levels appears as a major node in preserving microglial functions in the brain. Epigenetic mechanisms have been previously described to regulate *TREM2* gene expression and/or to be altered in the AD context. For instance, DNA methylation is increased in the promoter region of *TREM2* in AD brain samples compared to controls at relevant brain regions such as hippocampus (45) and prefrontal cortex (46–48). Non-coding RNAs also participate in the epigenetic modulation of gene expression. miRNA-34a has been reported to target *TREM2* mRNA 3’ untranslated region (UTR) in human sporadic AD hippocampal CA1 and in primary microglia cells cultures, so it appears that miRNA-34a is involved in the down-regulation of *TREM2* (56, 69, 70). In addition, the *TREM2* gene may be a source of epigenetic regulators, aimed at self-regulation or modulation of other genes expression.

Recently, circRNAs have emerged as interesting molecules that deserve to be investigated as epigenetic regulators and disease biomarkers. circRNAs are a particular type of non-coding RNAs that are generated by a process known as back-splicing from a pre-mRNA. As a consequence of back-splicing, circRNAs form covalently close loops which make them resistant to the action of RNase R and are, therefore, very stable molecules. circRNAs are thought to regulate alternative splicing and gene expression since back-splicing and linear splicing compete and both molecular processes share the spliceosome machinery (13, 26, 71).

Among the functions of circRNAs are regulation of parental gene expression by competition with its linear counterparts (9, 26), by modulating translation of cognate mRNA (13) or by occupying RNA binding sites in target genes (7). circRNAs can regulated protein expression and function by interacting with the protein or through protein complex (72). Another described function of circRNAs is the ability to translate into proteins as some of them have internal ribosomal entry site (IRES) within their sequences (32). circRNAs may also act as miRNAs sponges disrupting its function as inhibitors of mRNAs (28, 73).

The expression of circRNAs in the brain is high compared to other body tissues. Moreover, the expression of circRNAs in the brain changes throughout neuronal development, being in the adult phase when more circRNAs are detected; in other words, circRNAs accumulate with age. Their expression also changes according to brain region and cell subtypes. Research to date suggests that circRNAs may play important roles in synaptic plasticity and neuronal function (71, 74, 75).

Here, we describe a new circRNA transcript, circTREM2_1, arising from the microglial gene *TREM2,* in the human brain and microglial cells. To our knowledge, no other *TREM2*-derived circRNA has been previously reported. circTREM2_1 includes almost all the exons of *TREM2* gene (exons 2, 3, 4 and 5) including exon 4, which is relevant for *TREM2* functioning since encodes the transmembrane domain, along with most of cytoplasmatic and a small region of extracellular Trem2 protein. In addition, exon 4 skipping seems to participate in the synthesis of sTREM2, which is significantly increased in the CSF of AD patients. As circTREM2_1 includes exon 4, upregulation of circTREM2_1 may favour the increased expression of sTREM2. However, whether a relationship between sTREM2 and circTREM2_1 is present should be unravelled by further investigations.

In our study, we observed an upregulation of circTREM2_1 in the entorhinal cortex of AD compared to controls. The entorhinal cortex is a most vulnerable region where AD neuropathological changes start in AD. DAM have been described as associated with brain regions where AD changes are first shown. Thus, it would be very interesting to investigate whether increase in circTREM2_1 expression is detrimental in the case of neurodegenerative diseases, in particular in AD, and the role that circTREM2 may have in impairing DAM maintenance. Furthermore, increased expression levels of cirTREM2_1 were found mainly in the early phage of AD, suggesting that this circRNAs may be involved in the onset of the disease.

On the other hand, a negative correlation was shown between circTREM2_1 expression levels and Aβ deposits in the entorhinal cortex. This is in contrast with a positive correlation that was previously described for *TREM2* mRNA levels and hippocampal Aβ burden (45). Indeed, circTREM2_1 is elevated at the initial stage and then drops in intermediate and advanced stages. Whether circTREM2_1 has a regulatory role in *TREM2* mRNA expression should be analyzed by additional experiments. Furthermore, a number of miRNAs have been predicted to be linked to circTREM2s. The function of most of them is unknown at this time and represents area of special interest to be investigated.

To sum up, *TREM2* gene originates circTREM2_1, a novel circRNA which is upregulated in the entorhinal cortex of AD brains. Microglia is probably the main source of this circRNA in the human brain. Further investigations are needed to unravel the potential detrimental role of circTREM2_1 in the pathogenesis of AD.

## Supplementary Materials

Supplemental Figure 1, Supplemental Figure 2, Supplemental Figure 3, Supplemental Table 1 and Supplemental Table 2

## Acknowledgements

We want to kindly thank Isabel Gil-Aldea (Navarrabiomed Biobank, technical support) for their help. And we are very grateful to the patients and relatives that generously donate the brain tissue to the Navarrabiomed Biobank.

## Authors’ roles

AUC contributed to the study concept and design, running of the experiments, acquisition of the data, analysis and interpretation of the data, statistical analysis, figure drawing and drafting/revising of the manuscript for content. JSR contributed to the analysis and interpretation of the data (amyloid deposits), sorting of the patients into different stages, and acquisition of the image data. MRobles contributed to the running of the experiments and acquisition of the data. MRoldan contributed to the acquisition of the data. BA contributed to the acquisition of the data. MVZ participated in the revision of the subject diagnosis and classification of patients. IBL contributed to the study concept and design. MM contributed to the drafting/revising of the manuscript for content, study concept and design, analysis and interpretation of the data, study supervision, and obtaining the funding. All authors read and approved the final manuscript.

