## Supplementary figures and images for "A *TREM2*-derived circRNA is upregulated in the entorhinal cortex of Alzheimer’s disease patients"

### Supplemental Figure 1.jpg

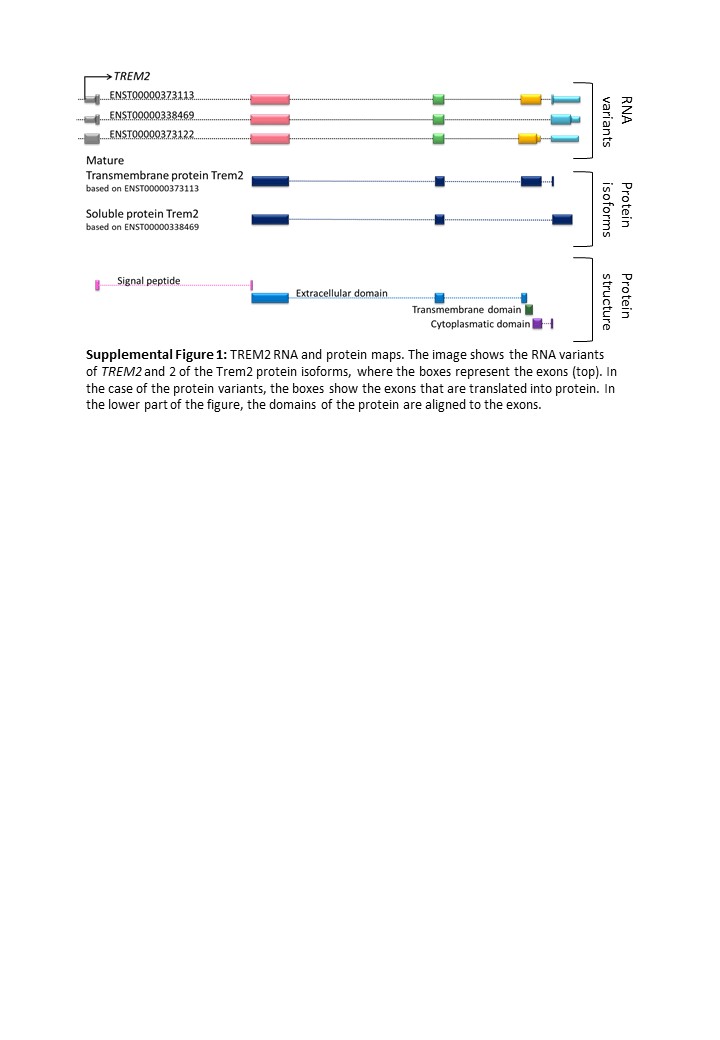

### Supplemental Figure 2.jpg

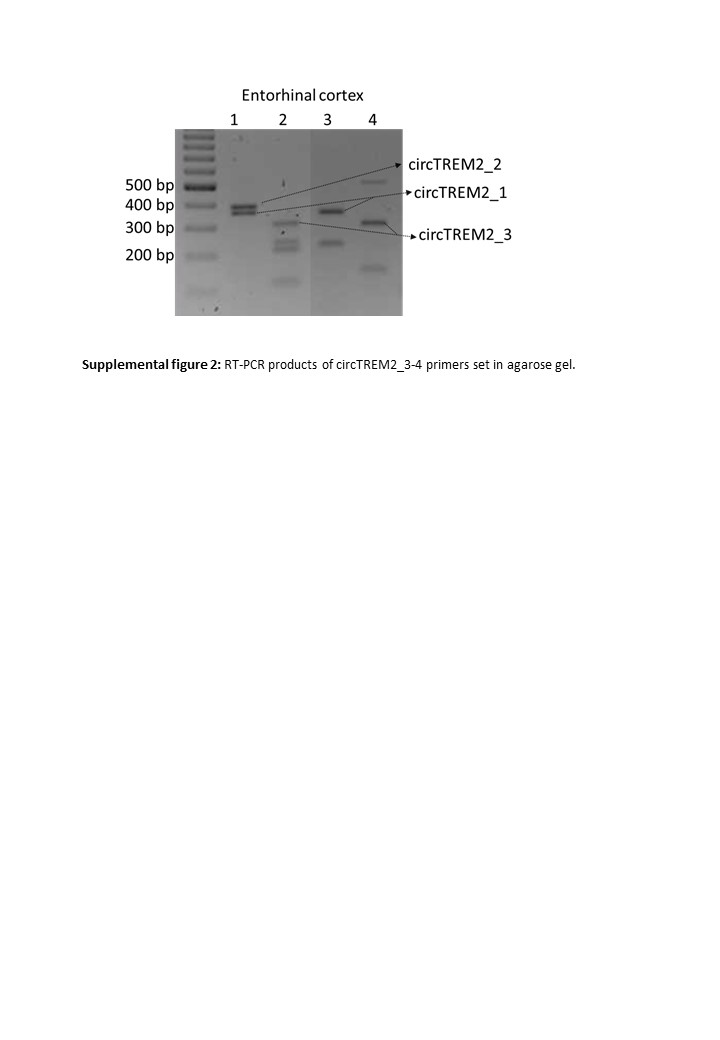

### Supplemental Figure 3.jpg

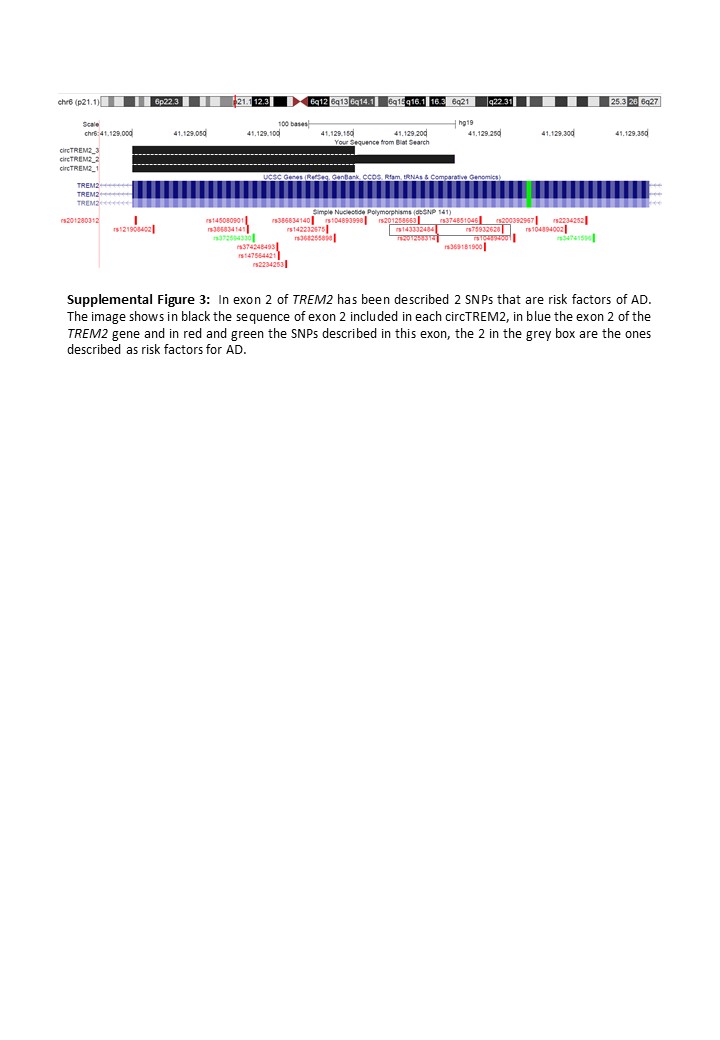
