## Supplemental Table 1 for "A *TREM2*-derived circRNA is upregulated in the entorhinal cortex of Alzheimer’s disease patients": Supplemental Table 1.pdf

**Supplemental Table 1. Brain sample set characteristics.** The table shows the characteristic of the samples included in the study. Nº: Number; NPD: no protein deposit; h: hours; PMI: *post mortem* interval.

| Nº | Braak stage | ABC score | ABC scale | Gender | % Aβ plaque area | Age at death (years) | PMI (h) |
|---|---|---|---|---|---|---|---|
| 1 | III | A1B2C1 | Low | Female | 0,349 | 96 | 1,5 |
| 2 | III | A1B2C1 | Low | Female | 0,044 | 88 | 33 |
| 3 | I | A1B1C1 | Low | Female | 0.00 | 85 | 4,3 |
| 4 | NA | NA | NA | Female | 0,672 | 69 | NA |
| 5 | II | A1B1C1 | Low | Female | 0,147 | 66 | 1,4 |
| 6 | III | A1B2C3 | Int | Female | 0,648 | 84 | 13 |
| 7 | IV | A2B2C2 | Int | Female | 0,814 | 97 | NA |
| 8 | IV | A2B2C3 | Int | Male | 0,420 | 78 | 5 |
| 9 | I | A1B1C1 | Low | Male | 0,000 | 60 | 15,3 |
| 10 | V | A3B3C2 | High | Male | NA | 91 | 5 |
| 11 | III | A3B2C3 | Int | Male | 0,724 | 83 | 9 |
| 12 | IV | A3B2C1 | Int | Female | 0,131 | 90 | 3 |
| 13 | I | A1B1C1 | Low | Male | 0,024 | 85 | 3,2 |
| 14 | III | A3B2C3 | Int | Female | 0,376 | 85 | NA |
| 15 | III-IV | A3B2C3 | Int | Female | 0,357 | 98 | 3 |
| 16 | IV | A2B2C2 | Int | Female | 0,431 | 91 | 10 |
| 17 | III | A2B2C3 | Int | Female | 0,528 | 98 | 23 |
| 18 | V | A3B3C3 | High | Female | 0,777 | 77 | 11 |
| 19 | VI | A3B3C3 | High | Female | 0,777 | 86 | 2,3 |
| 20 | V | A3B3C2 | High | Female | 1,610 | 82 | 9 |
| 21 | I | A1B1C1 | Low | Female | NA | 85 | 2 |
| 22 | IV | A2B2C2 | Int | Male | 0,959 | 88 | 3,5 |
| 23 | VI | A3B3C3 | High | Male | 2,013 | 70 | 2,35 |
| 24 | II | A2B1C3 | Low | Male | NA | 80 | 3 |
| 25 | II | A1B1C1 | Low | Male | NA | 74 | 2,5 |
| 26 | II | A1B2C2 | Int | Female | NA | 71 | 4 |
| 27 | II | A2B1C1 | Low | Female | NA | 80 | 3,7 |
| 28 | 0 | control | Not | Female | NPD | 43 | 3 |
| 29 | 0 | control | Not | Male | NPD | 54 | 18 |
| 30 | 0 | control | Not | Female | NPD | 19 | NA |
| 31 | 0 | control | Not | Female | NPD | 46 | 7 |
| 32 | 0 | control | Not | Male | NPD | 28 | 6 |
| 33 | 0 | control | Not | Male | NPD | 41 | 3,5 |
| 34 | 0 | control | Not | Male | NPD | 54 | 2,7 |
| 35 | 0 | control | Not | Male | NPD | 81 | 10,5 |
| 36 | 0 | control | Not | Male | NPD | 26 | 6,2 |
| 37 | 0 | control | Not | Male | NPD | 53 | 7 |
| 38 | 0 | control | Not | Female | NPD | 88 | 9 |
| 39 | 0 | control | Not | Male | NPD | 66 | 6,5 |
| 40 | 0 | control | Not | Female | NPD | 88 | 3,5 |
| 41 | 0 | control | Not | Female | NPD | 76 | 11,5 |
| 42 | 0 | control | Not | Male | NPD | 65 | 3 |
| 43 | 0 | control | Not | Male | NPD | 83 | NA |
