## Supplemental table 2 for "A *TREM2*-derived circRNA is upregulated in the entorhinal cortex of Alzheimer’s disease patients": Supplemental Table 2.pdf

| | | Forward 5'->3' | Reverse 5'->3' | Amplicon molecular weight (bp) expected for each circTREM2/primer set: circTREM2_1 | circTREM2_2 | circTREM2_3 |
|---|---|---|---|---|---|---|
| **divergent PCR primers set** | **circTREM2_4-5** | GTGGGGAGGTGGTAAGAACA | CCAGGAGGAGAAGGATGGA | - | - | - |
| | **circTREM2_3-4** | ACAGAAGCCAGGGACACATC | CATCCTCGAAGCTCTCAGAC | 358 | 427 | 306 |
| | **circTREM2_2-4** | ACAGAAGCCAGGGACACATC | CTGGTAGAGACCCGCATCAT | 228 | 297 | 176 |
| | **circTREM2_2-3** | CATGTGGAGCACAGCATCTC | CTGGTAGAGACCCGCATCAT | 368 | 437 | 316 |
| | **circTREM2_3-5** | GAAAAGCCCAGGAGAAGTCC | GGAACCAGAGATCTCCAGCA | 221 | - | 290 |

| | | Forward 5'->3' | Reverse 5'->3' | Probe 5'->3' |
|---|---|---|---|---|
| **TaqMan assay** | **circTREM2** | CAGGGTATCAGCTCCAAACTC | AGCGTAATGGTGAGAGTGC | ATCGTCTGTGATGGCTGTGCTGG |
| | **ACTB** | ACCTTCTACAATGAGCTGCG | CCTGGATAGCAACGTACATGG | ATCTGGGTCATCTTCTCGCGGTTG |

**Supplemental Table 2:** Sequence of set of primers used for RT-PCR and RTqPCR experiments.
